# Genetic contribution to ‘theory of mind’ in adolescence

**DOI:** 10.1101/254524

**Authors:** Varun Warrier, Simon Baron-Cohen

## Abstract

Difficulties in ‘theory of mind’ (the ability to attribute mental states to oneself or others, and to make predictions about another’s behaviour based on these attributions) have been observed in several psychiatric conditions. We investigate the genetic architecture of theory of mind in 4,577 13-year-olds who completed the Emotional Triangles Task (Triangles Task), a first-order test of theory of mind. We observe a small but significant female-advantage on the Triangles Task (Cohen’s d = 0.19, P < 0.01), in keeping with previous work using other tests of theory of mind. Genome-wide association analyses did not identify any significant loci, and SNP heritability was small and non-significant. Polygenic scores for six psychiatric conditions (ADHD, anorexia, autism, bipolar disorder, depression, and schizophrenia), and empathy were not associated with scores on the Triangles Task. However, polygenic scores of cognitive aptitude, and cognitive empathy, a term synonymous with theory of mind and measured using the “Reading the Mind in the Eyes” Test, were significantly associated with scores on the Triangles Task at multiple P-value thresholds, suggesting shared genetics between different measures of theory of mind and cognition.

## Introduction

Theory of mind is the ability to attribute mental states to one self and others and to use such mental state attribution to make sense of behaviour and predict it. First order theory of mind refers to the ability to understand another person’s mental state (e.g., “He thinks x”). Second-order theory of mind is when theory of mind is applied recursively to understand what someone is thinking of another person’s mental state (e.g., “He thinks that she thinks x”). Typically, first order theory of mind develops in early childhood (by 3 to 4 years of age)^1^, though precursors to theory of mind are evident at the end of infancy, around 9-14 months of age, in joint attention behaviours such as proto-declarative pointing and gaze following^2^. This suggests a developmental component to theory of mind, including a social learning component. Second-order theory of mind develops a little later, by age 5 to 6 years of age^3^. Other studies have identified that infants show an implicit understanding other’s mental states using looking-time towards a location where another person believes an object will be^4^. The development of theory of mind largely follows consistent patterns, irrespective of culture, suggests it is a human universal that might have evolved and thus be partly heritable^5^.

Due to the complex nature of theory of mind, twin studies have identified different heritabilities for theory of mind and related phenotypes at different developmental stages. Heritability is also different for first-order and second-order theory of mind tasks. No study has investigated the twin heritability of the task used in this study – the Emotional Triangles Task (Triangles Task), a first-order test of theory of mind^6^. However, a few studies have investigated the twin heritability of other theory of mind tasks. A large study investigating the heritability of different theory of mind tasks in 1,116 5-year olds, and suggested that shared environmental influences rather than genetic factors contribute to most of the variance in these tasks^7^. Another study in 695 9-year-olds identified a small, but non-significant additive genetic component^8^. However, other studies have identified modest heritabilities in theory of mind and related phenotypes. A study based on parent-reports of children’s prosocial and antisocial behaviour requiring theory of mind in 2 – 4 year olds identified a modest and significant heritability^9^. In adults, cognitive empathy, measured using the ‘Reading the Mind in the Eyes’ Test (Eyes Test), identified a significant twin heritability of approximately 28%^10^. The term ‘cognitive empathy’ is synonymous with theory of mind, although in the Eyes test includes a visual recognition element of another’s mental state, including their emotion. In contrast, ‘affective empathy’ is the ability to respond appropriately to another’s mental state. Together, cognitive and affective empathy are two major facets of empathy^11,12^.

Difficulties in theory of mind have been identified in different psychiatric conditions. Children with autism have difficulties in attributing mental states^13^, known as ‘mindblindness’. This comes by degrees, rather than being all or none. This may be manifested in children with autism developing theory of mind abilities later than age and IQ matched typical controls. Adults with autism also show difficulties in theory of mind, using age-appropriate tasks^14,15^. Similarly, a meta-analysis identified significant impairments in tasks involving theory of mind in individuals with schizophrenia^16^. Difficulties in theory of mind have also been identified in people with unipolar and bipolar disorders^17–19^, eating disorders^20^, and attention deficit and hyperactivity disorder or ADHD^21^. For example, a recent meta-analysis of theory of mind in people with eating disorders (15 studies, 677 cases and 514 controls) identified significant deficits in theory of mind in individuals with anorexia compared to typical controls^22^. In ADHD, theory of mind difficulties are associated with deficits in executive function^23^. Theory of mind is also predicted by measures of general cognition such as IQ and working memory^24,25^.

These differences in performance on tests of theory of mind could be due to underlying biology, or other environmental processes that mediate performance on tests of theory of mind in individuals with psychiatric conditions. Here, we test the genetic correlates of first-order theory of mind using the Emotional Triangles Task (Triangles Task) ^6^. In the Triangles Task, participants are required to attribute mental states to animated triangles (e.g., “the triangle is angry”). In the original version of this task, the participant is simply asked to describe what they see, and the spontaneous narratives are coded for the number and type of mental state attribution. In the version used in the current study, participants are asked to pick the right mental state (from one of four options in a forced choice format) based on motion-cues of the triangles. The sample comprised 4,577 13 year olds from the Avon Longitudinal Study of Parents and Children on whom we had both genetic and phenotypic data, after quality control. They took the Emotional Triangles task during adolescence, a period marked by key changes in neural architecture, in peer-relationship, and in hormonal profile^26,27^. Interrogating the genetic relationship between theory of mind at this age and risk for psychiatric conditions with known difficulties in theory of mind (autism, ADHD, anorexia nervosa, bipolar disorder, depression, and schizophrenia) may identify genetic biomarkers.

This study has three specific aims: 1. To determine the narrow-sense heritability (or SNP-chip heritability) of theory of mind in 13-year-olds, measured using the Emotional Triangles Task; 2. To identify any genes and genetic loci associated with the Triangles Task; and 3. To test if polygenic risk scores for six psychiatric conditions (ADHD, anorexia, autism, bipolar disorder, depression and schizophrenia), cognitive aptitude, and two different measures of empathy (the EQ^28^ and the Eyes Test^10^) predict performance on the Triangles Task in 13-year-olds.

## Methods

### Phenotype and participants

Theory of mind was measured using the Emotional Triangles Task (Triangles Task)^6^. All participants were 13 years of age (born in April 1991- Dec 1992), and measures were collected as a part of the ongoing Avon Longitudinal Study of Parents and Children (ALSPAC). Data was queried using the fully searchable data dictionary, which is available online at http://www.bristol.ac.uk/alspac/researchers/access/. ALSPAC consists of 14,541 initial pregnancies from women resident in Avon, UK resulting in a total of 13,988 children who were alive at 1 year of age. In addition, children were enrolled in additional phases, which are described in greater detail elsewhere^29^.

The study received ethical approval from the ALSPAC Ethics and Law Committee, and written informed consent was obtained from parent or a responsible legal guardian for the child to participate. Assent was obtained from the child participants where possible. In addition, we also received ethical permission to use de-identified summary genetic and phenotype data from the Human Biology Research Ethics Committee at the University of Cambridge. All research was performed in accordance to the Helsinki Declaration.

The Triangles Task is a test of theory of mind where participants have to attribute mental states to non-living shapes (animated triangles). Test-retest reliability of the mental state coding scheme has identified an interclass correlation of 0.69 and a technical error of measurement of 0.66^6^. The Triangles Task consists of 28 questions (16 scored questions and 12 control questions). In each question, a 5-second animation of a triangle is shown and a question is asked about the mental state of the triangle (e.g. Was the triangle angry?). Participants are asked to choose from 0 to 5 (a Likert-scale) to respond to the question, where 0 indicates that the triangle did not possess the mental state described in the question, and 5 indicates that the triangle definitely possessed the mental state described in the question. In total, four mental states were tested (happy, sad, angry, and scared). For each mental state, there were two positive questions, where the mental state of the triangle matched the mental state described in the question, and two negative questions, where the mental state of the triangle did not match the mental state described in the question (e.g. the triangle is shown to be happy, and the subsequent question is “Was the triangle sad?”). Hence, in total, there were 16 questions that were scored. Control questions comprised of asking if the triangle was living, and participants, again, had to choose between 0 - 5.

We calculated the total score by adding the score for all the positive questions and subtracting the score for the negative items. To avoid negative scores, we added 40 to the total score, giving the score a range from 0 - 80. We removed participants on whom we did not have any genetic data. We removed participants who had chosen the same answer for more than 50% of the items, including the control items, suggesting that they were not attending to the task. This was done after carefully evaluating the options selected and the reaction times. We noticed that, for these participants that 1. The reaction time was small for three or more consecutive items at various points in the test 2; the same option was chosen for three or more consecutive items at various points on the tests; 3. the same option was chosen for both the positive and negative questions; and 4. The control questions were answered incorrectly at various points in the test. After removing these participants, we had phenotypic and genetic data on 4,577 participants (n = 2,217 females, and n = 2,360 males).

### Genotyping and Imputation

Genotyping and imputation was conducted by ALSPAC. All participants were genotyped using the Illumina^®^ HumanHap550 quad chip by 23andMe. GWAS data was generated by Sample Logistics and Genotyping Facilities at Wellcome Sanger Institute and LabCorp (Laboratory Corportation of America), and with support from personal genomics company 23andMe., Inc. This resulting raw genome-wide data were subjected to the following quality control procedures: Individuals were removed with discordant gender information, if there was excessive or low genetic heterozygosity, if missingness was > 3%, if they were of non-European ancestry as measured using multidimensional scaling analyses compared with Hapmap II (release 22), and if there was evidence of cryptic relatedness (>10% IBD). SNPs were removed if they had a minor allele frequency < 1%, deviated from Hardy-Weinberg equilibrium (P < 5x10^-7^), and had a call rate < 95%. This resulted in a total of 526,688 genotyped SNPs. Using SNP data from mothers and children (477,482 common SNPs between mothers and children), haplotypes were estimated using ShapeIT (v2.r644)^30^. Imputation was performed using Impute2 V2.2^31^ against the 1000 genomes reference panel (Phase 1, Version 3), using all 2186 reference haplotypes including non-Europeans. Imputed SNPs were excluded from all further analyses if they had a minor allele frequency < 1% and info < 0.8, which resulted in a total of 8,282,911 SNPs.

### Genome-wide association analyses and gene based analyses

We conducted a genome-wide association analyses (GWAS) on the total score on the Triangles Task. In addition to the quality control procedure described above, we conducted additional quality control steps for the participants included in this study. We included only those SNPs with a minor-allele frequency > 1%. We excluded all SNPs that were not in Hardy-Weinberg equilibrium (P < 5 x 10^-7^), had a per-SNP missing rate > 5%. Similarly, we excluded all individuals who had a genotype missing rate > 10%. We included sex and the first two genetic ancestry principal components. Regression analyses was run using a linear regression model in Plink 1.9 ^32^. BGEN files were converted to Plink format using hard calls. Calls with uncertainty greater than 0.1 were treated as missing. After quality control, 4,577 were included in the GWAS. Gene-based analyses was conducted using MAGMA^33^. Significant genes were identified after Bonferroni correction (P < 2.74x10^-6^).

### Heritability and Polygenic risk scores

SNP heritability was estimated using GCTA v1.26^34^ (GREML) and LDSC^35^. For GCTA GREML, heritability was calculated after including sex and the first two genetic principal components as covariates. We investigated for inflation in Chi-square statistics due to uncorrected population stratification using LDSC^35^. Given that the cohort was comprised of unrelated individuals and individuals with cryptic relatedness were removed (IBD > 10%) during quality control of the raw genotype data, we calculated the genetic relatedness matrix using all individuals in the study.

Given the different polygenicity and power of the GWAS used as the training datasets, we constructed polygenic risk score using PRSice^36^ at eight different P-value thresholds (0.01, 0.05, 0.1, 0.15, 0.20, 0.5, 0.8 and 1.0). We chose these thresholds so as to balance the signal to noise ratio in GWAS results used as training datasets. PRSice calculates a weighted-average score of all the relevant alleles. Clumping was performed in PRSice using an r^2^ of 0.1. We used a linear model to regress the polygenic scores and the covariates against the scores on the Triangles Task. Polygenic risk scores were constructed using summary data for 6 psychiatric conditions (ADHD^37^ (n = 55,374), anorexia^38^ (n = 14,477), autism^39^ (n = 15,954), bipolar disorder^40^ (n = 16,731), major depressive disorder^41^ (n = 16,610), and schizophrenia^42^ (n = 79,845), cognitive aptitude^43^ (n = 78,803), self-reported empathy (EQ^28^) (n = 46,861), and cognitive empathy (the Eyes Test)^10^ (n = 89,553). We excluded educational attainment from the current analyses given that the summary GWAS data available for the educational attainment GWAS included participants from the ALSPAC and is likely to increase the probability of false positives. We included sex, and the first two ancestry principal components as covariates for polygenic scoring. We did not include age as all participants were tested at approximately 13 years of age. We used a Benjamini Hochberg FDR correction to correct for the tests conducted.

### Data availability

Data were downloaded from the Psychiatric Genomics Consortium (PGC) website for the 6 psychiatric conditions (http://www.med.unc.edu/pgc/results-and-downloads), and the Complex Traits Genomics website for cognitive aptitude (https://ctg.cncr.nl/software/summary_statistics). Data for the EQ and the Eyes Test were obtained from 23andMe, Inc.

## Results

### Phenotypic distribution

The range of the scores of the participants was 28 - 80. Inspection of the frequency histogram and quantile-quantile plot suggested a normal distribution **(Figure 1)**. The mean score of the participants was 56.93 (SD = 7.43). Females scored significantly higher than males (Females: 57.68, SD = 7.43; Males: 56.22, SD = 7.36; P < 0.001, unpaired, two-tailed T-test), though the effect size was small (Cohen’s d = 0.19).

**Figure 1:**
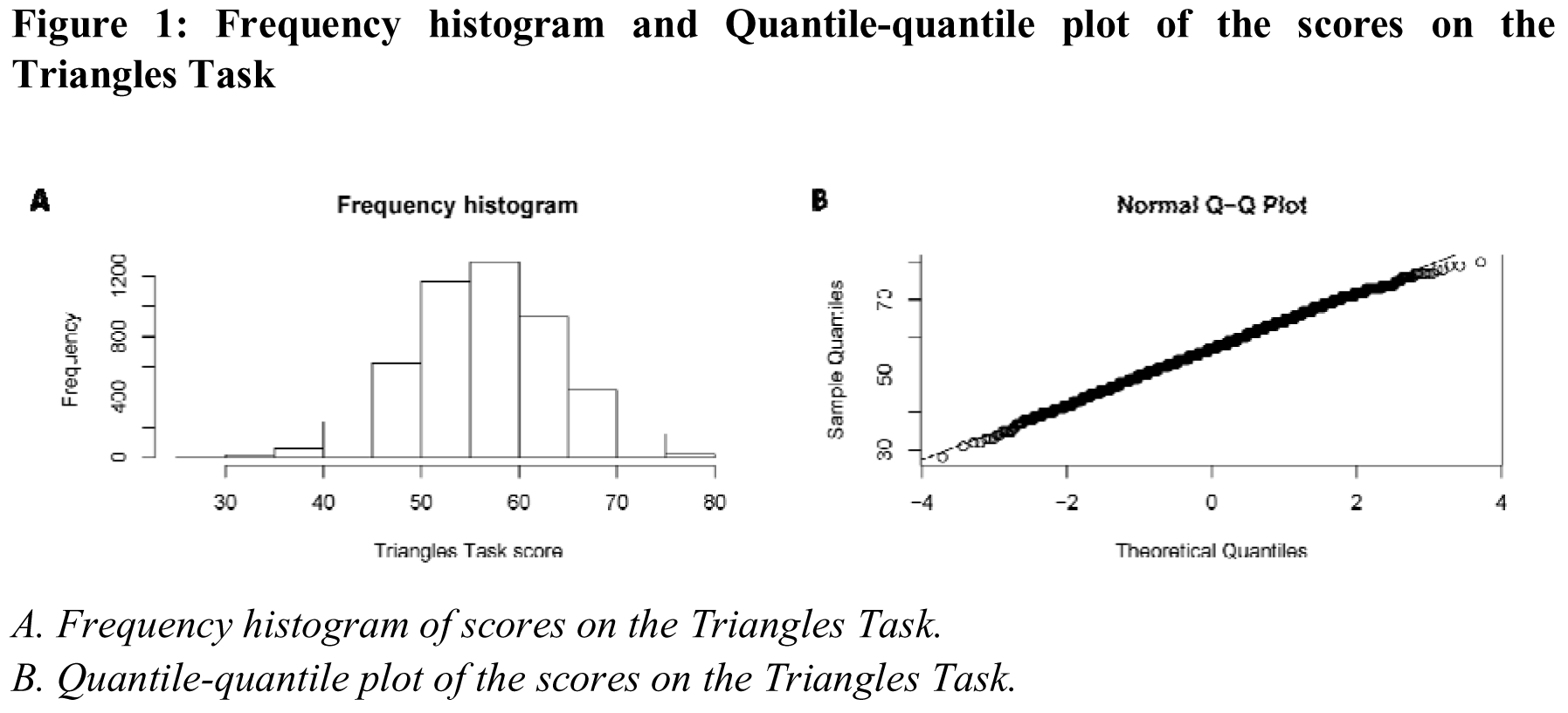
Frequency histogram and Quantile-quantile plot of the scores on the Triangles Task

### Genome-wide association analyses and heritability

Genome-wide association analyses did not identify any significant SNP. The sentinel SNP (SNP with the lowest P-value at the locus) at the most significant locus was rs2120452 (P = 6.8x10^-7^) on chromosome 1. The SNP lies in a non-coding RNA LOC105372904. The sentinel SNP at the second-most significant locus was rs17753687 (P = 8.6x10^-7^), which is an intronic SNP in *BBS4*, a gene implicated in Bardet–Biedl syndrome. Investigation of the QQ plots and LD score regression intercept did not reveal any inflation in effect sizes due to population stratification (LD score regression intercept = 0.99 ±0.0063). SNP heritability was small and nonsignificant (LDSR - h^2^_SNP_ = 0.13 ± 0.10; P = 0.16; GCTA - h^2^_SNP_ = 0.072±0.069; P = 0.29). Manhattan and QQ-plots are provided in **Figure 2.**

**Figure 2:**
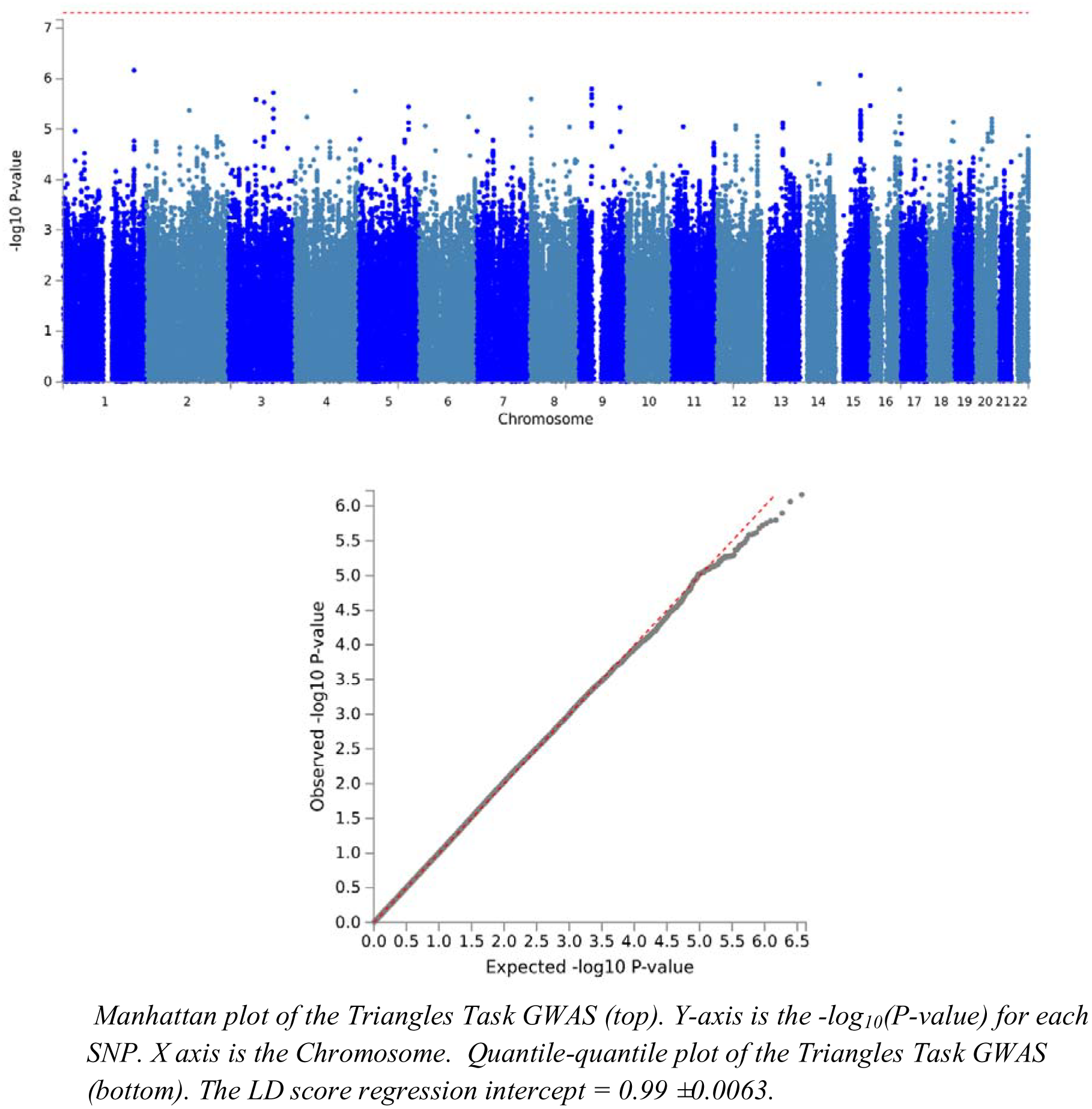
Manhattan plot and quantile-quantile plot of the GWAS of the Triangles Task

### Gene-based analysis

We conducted gene-based analysis using MAGMA, and did not identify and significant genes at a genome-wide P-value threshold of 2.74x10^-6^. The most significant gene was *MARK4* at 19q13.32 (P = 2.96x10^-6^). MARK4 is involved in phosphorylating microtubule associated proteins. It has high expression in the brain and in testes, according to GTEx^44^.

### Polygenic risk scores

As the heritability was non-significant to conduct genetic correlation analyses, we conducted polygenic score regression with 6 psychiatric conditions, cognitive aptitude, cognitive empathy, and self-reported empathy. We tested polygenic score at 8 P-value thresholds. We did not identify a significant polygenic score after FDR-based correction for any of the six psychiatric conditions investigated **(Figure 3)**. Overall, the polygenic scores predicted limited variance for psychiatric conditions **(Table 1)**. However, polygenic scores in both cognitive aptitude and cognitive empathy were significantly associated with scores in the Triangles Task across all the thresholds tested using after FDR correction. For cognitive aptitude and at two P-value thresholds for cognitive empathy, these results were significant even after using a more stringent Bonferroni correction **(Figure 3 and Table 1)**. This likely reflects both the greater statistical power of the two datasets when compared to the other GWAS datasets in the condition and the underlying pleiotropy between theory of mind and cognition as previous studies have identified a modest, positive correlation between different measures of theory of mind and cognition ^10,45^.

**Figure 3:**
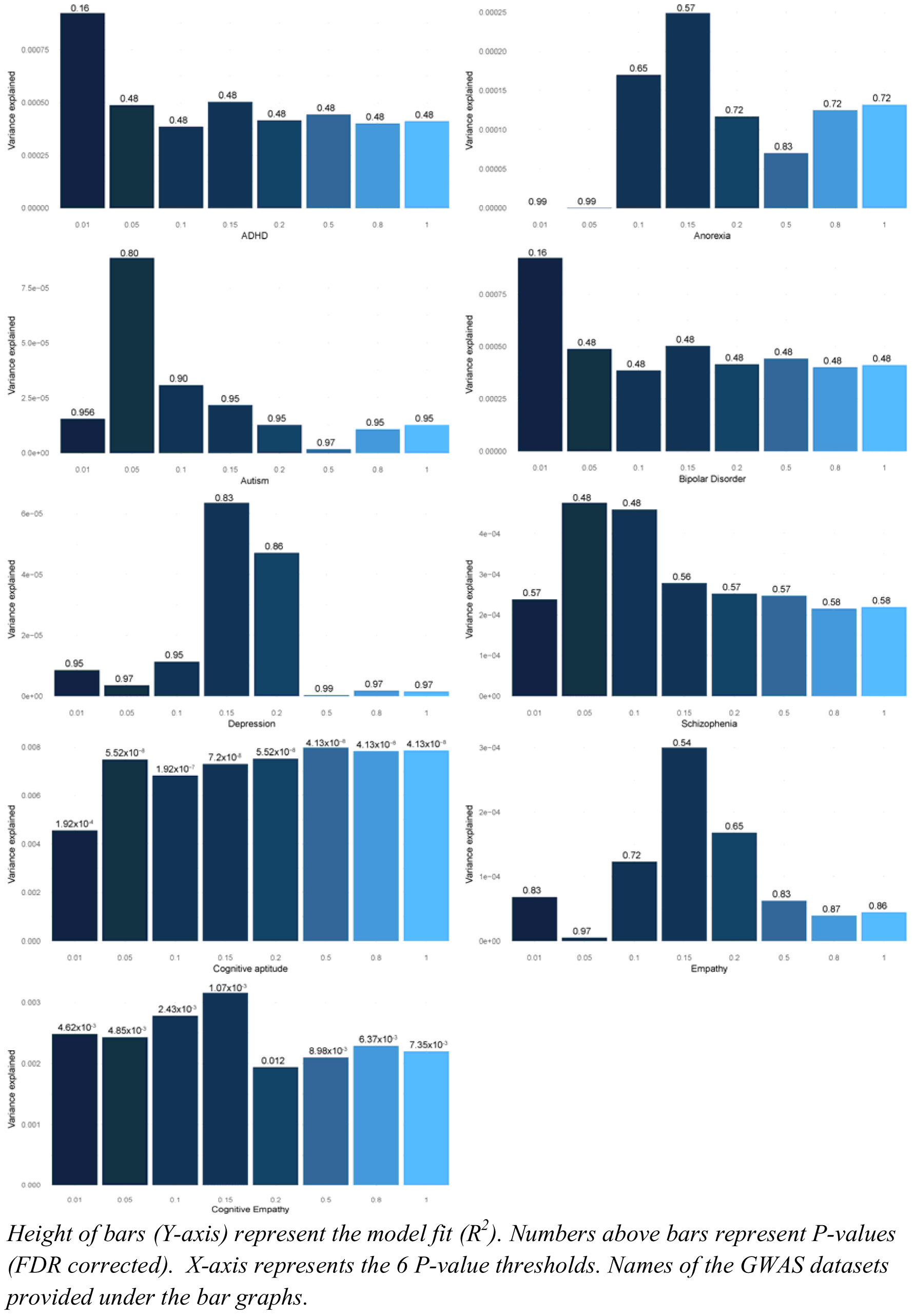
Polygenic score results at various P-value thresholds for the Triangles Task

**Table 1:** Results of the Polygenic score analyses

| Threshold | P | r<sup>2</sup> | FDR P | Phenotype |
| --- | --- | --- | --- | --- |
| 0.01 | 9.94E-01 | 1.37E-08 | 9.94E-01 | Anorexia |
| 0.05 | 9.90E-01 | 3.40E-08 | 9.94E-01 | Anorexia |
| 0.1 | 3.75E-01 | 1.70E-04 | 6.50E-01 | Anorexia |
| 0.15 | 2.83E-01 | 2.49E-04 | 5.72E-01 | Anorexia |
| 0.2 | 4.63E-01 | 1.17E-04 | 7.25E-01 | Anorexia |
| 0.5 | 5.69E-01 | 7.03E-05 | 8.34E-01 | Anorexia |
| 0.8 | 4.47E-01 | 1.25E-04 | 7.22E-01 | Anorexia |
| 1 | 4.36E-01 | 1.32E-04 | 7.22E-01 | Anorexia |
| 0.01 | 7.90E-01 | 1.54E-05 | 9.56E-01 | Autism |
| 0.05 | 5.23E-01 | 8.86E-05 | 8.01E-01 | Autism |
| 0.1 | 7.06E-01 | 3.08E-05 | 9.08E-01 | Autism |
| 0.15 | 7.52E-01 | 2.17E-05 | 9.50E-01 | Autism |
| 0.2 | 8.08E-01 | 1.27E-05 | 9.56E-01 | Autism |
| 0.5 | 9.30E-01 | 1.69E-06 | 9.74E-01 | Autism |
| 0.8 | 8.23E-01 | 1.08E-05 | 9.56E-01 | Autism |
| 1 | 8.08E-01 | 1.27E-05 | 9.56E-01 | Autism |
| 0.01 | 3.90E-02 | 9.24E-04 | 1.65E-01 | ADHD |
| 0.05 | 1.33E-01 | 4.88E-04 | 4.86E-01 | ADHD |
| 0.1 | 1.83E-01 | 3.85E-04 | 4.86E-01 | ADHD |
| 0.15 | 1.28E-01 | 5.03E-04 | 4.86E-01 | ADHD |
| 0.2 | 1.66E-01 | 4.15E-04 | 4.86E-01 | ADHD |
| 0.5 | 1.53E-01 | 4.43E-04 | 4.86E-01 | ADHD |
| 0.8 | 1.74E-01 | 4.01E-04 | 4.86E-01 | ADHD |
| 1 | 1.69E-01 | 4.11E-04 | 4.86E-01 | ADHD |
| 0.01 | 8.52E-01 | 7.54E-06 | 9.59E-01 | Bipolar disorder |
| 0.05 | 6.31E-01 | 5.01E-05 | 8.68E-01 | Bipolar disorder |
| 0.1 | 3.68E-01 | 1.76E-04 | 6.50E-01 | Bipolar disorder |
| 0.15 | 2.43E-01 | 2.96E-04 | 5.47E-01 | Bipolar disorder |
| 0.2 | 2.23E-01 | 3.23E-04 | 5.35E-01 | Bipolar disorder |
| 0.5 | 1.89E-01 | 3.73E-04 | 4.86E-01 | Bipolar disorder |
| 0.8 | 1.86E-01 | 3.79E-04 | 4.86E-01 | Bipolar disorder |
| 1 | 1.99E-01 | 3.57E-04 | 4.94E-01 | Bipolar disorder |
| 0.01 | 8.42E-01 | 8.59E-06 | 9.59E-01 | Depression |
| 0.05 | 8.98E-01 | 3.56E-06 | 9.74E-01 | Depression |
| 0.1 | 8.20E-01 | 1.13E-05 | 9.56E-01 | Depression |
| 0.15 | 5.88E-01 | 6.36E-05 | 8.34E-01 | Depression |
| 0.2 | 6.41E-01 | 4.71E-05 | 8.68E-01 | Depression |
| 0.5 | 9.71E-01 | 2.88E-07 | 9.94E-01 | Depression |
| 0.8 | 9.28E-01 | 1.76E-06 | 9.74E-01 | Depression |
| 1 | 9.33E-01 | 1.51E-06 | 9.74E-01 | Depression |
| 0.01 | 2.95E-01 | 2.38E-04 | 5.74E-01 | Schizophrenia |
| 0.05 | 1.38E-01 | 4.76E-04 | 4.86E-01 | Schizophrenia |
| 0.1 | 1.46E-01 | 4.59E-04 | 4.86E-01 | Schizophrenia |
| 0.15 | 2.57E-01 | 2.78E-04 | 5.61E-01 | Schizophrenia |
| 0.2 | 2.81E-01 | 2.52E-04 | 5.72E-01 | Schizophrenia |
| 0.5 | 2.86E-01 | 2.47E-04 | 5.72E-01 | Schizophrenia |
| 0.8 | 3.19E-01 | 2.15E-04 | 5.89E-01 | Schizophrenia |
| 1 | 3.15E-01 | 2.19E-04 | 5.89E-01 | Schizophrenia |
| 0.01 | 7.06E-04 | 2.48E-03 | 4.62E-03 | Cognitive empathy |
| 0.05 | 8.08E-04 | 2.43E-03 | 4.85E-03 | Cognitive empathy |
| 0.1 | 3.38E-04* | 2.78E-03 | 2.43E-03 | Cognitive empathy |
| 0.15 | 1.34E-04* | 3.16E-03 | 1.07E-03 | Cognitive empathy |
| 0.2 | 2.78E-03 | 1.94E-03 | 1.25E-02 | Cognitive empathy |
| 0.5 | 1.87E-03 | 2.10E-03 | 8.98E-03 | Cognitive empathy |
| 0.8 | 1.15E-03 | 2.29E-03 | 6.37E-03 | Cognitive empathy |
| 1 | 1.43E-03 | 2.20E-03 | 7.35E-03 | Cognitive empathy |
| 0.01 | 5.75E-01 | 6.82E-05 | 8.34E-01 | Empathy |
| 0.05 | 8.79E-01 | 5.01E-06 | 9.74E-01 | Empathy |
| 0.1 | 4.51E-01 | 1.23E-04 | 7.22E-01 | Empathy |
| 0.15 | 2.39E-01 | 3.00E-04 | 5.47E-01 | Empathy |
| 0.2 | 3.79E-01 | 1.68E-04 | 6.50E-01 | Empathy |
| 0.5 | 5.91E-01 | 6.27E-05 | 8.34E-01 | Empathy |
| 0.8 | 6.69E-01 | 3.96E-05 | 8.76E-01 | Empathy |
| 1 | 6.51E-01 | 4.44E-05 | 8.68E-01 | Empathy |
| 0.01 | 2.13E-05* | 4.56E-03 | 1.92E-04 | Cognitive aptitude |
| 0.05 | 3.83E-09* | 7.50E-03 | 5.52E-08 | Cognitive aptitude |
| 0.1 | 1.87E-08* | 6.83E-03 | 1.92E-07 | Cognitive aptitude |
| 0.15 | 6.00E-09* | 7.31E-03 | 7.20E-08 | Cognitive aptitude |
| 0.2 | 3.54E-09* | 7.53E-03 | 5.52E-08 | Cognitive aptitude |
| 0.5 | 1.21E-09* | 7.98E-03 | 4.13E-08 | Cognitive aptitude |
| 0.8 | 1.72E-09* | 7.83E-03 | 4.13E-08 | Cognitive aptitude |
| 1 | 1.61E-09* | 7.86E-03 | 4.13E-08 | Cognitive aptitude |

*\*indicates significant polygenic score association after Bonferroni correction ( $P < 6.94 \times 10^{-4}$ )*

## Discussion

We investigated the genetic correlates of first-order theory of mind using the Triangles Task. In total, 4,577 13-year-olds completed the Triangles Task, making this the largest investigation of genetic correlates of theory of mind at a specific age. At a phenotypic level, the scores on the Triangles Task were normally distributed and we observed a small but significant female-advantage on the Triangles Task. This is similar to what has been observed in other studies of cognitive empathy ^46^ and facial expression recognition^47^.

The current study finds limited evidence for a genetic contribution to the Triangles Task in 13-year-olds in this sample. Genome-wide association analyses did not identify any significant loci at P < 5x10^-8^. Furthermore, gene-based analyses also did not identify any significant genes. We note, however, that the current study is statistically underpowered. Previous work on the genetics of cognitive empathy, which is related to theory of mind had identified that the per-SNP variance explained for the most significant SNP was 0.013% after correcting for winner’s curse ^10^. Post-hoc power calculations suggest that a sample two orders of magnitude larger than the current sample would be required to identify genome-wide significant loci, if the effect sizes are similar. This, however, is challenging given the nature of the task, which demands that participants spend at least half an hour to complete the task. We also note that we are statistically underpowered to identify significant additive SNP heritability, assuming a true additive SNP heritability of 5%, which is similar to the SNP heritability reported elsewhere^10^. These calculations preclude us from conducting genetic correlation analyses using the current cohort.

We also investigated if polygenic scores from 6 psychiatric conditions, empathy, cognitive empathy and cognitive aptitude predict performance on the Triangles Task. We used PRSice and investigated the predictive power of polygenic scores at eight different P-value thresholds providing reasonable resolution. We note that the sample sizes for the training GWAS set are varied, although all the GWAS datasets had more than 10,000 participants. However, polygenic scores for none of the psychiatric conditions significantly predicted performance on the Triangles Task across the six different P-value thresholds. In contrast, polygenic scores for cognitive empathy as measured using the Eyes Test, and cognitive aptitude significantly predicted variance in the Triangles Task, underscoring previously observed results^10^.

Our results indicate that genetic risk for psychiatric conditions do not explain much of the variance in theory of mind ability in adolescents in this sample. We speculate that this must be due to different reasons. First, it is likely that the current task does not capture the entire variance in theory of mind. Indeed, as mentioned earlier, theory of mind is complex and designing a task to capture the intrinsic variance in theory of mind is challenging. The Triangles task only considers first order mental state attributions, and the range of mental states is very limited, compared to for example the ‘Reading the Mind in the Eyes’ test^14^. It is also likely that the difficulties in theory of mind observed in individuals with psychiatric conditions may be due to other processes that mediate theory of mind. Interrogation of the genetic architecture of diverse phenotypes that contribute to social behaviour and theory of mind will help understand how they contribute to genetic risk for various psychiatric conditions. We can also not exclude the possibility that using either a larger training dataset and/or a larger target dataset will help improve the statistical significance of the polygenic score association. Finally, we cannot ignore non-genetic contributors to theory of mind. Certainly, twin studies do suggest that for certain theory of mind tasks, the genetic contribution is negligible.

Previous work from our lab investigated the genetic architecture of cognitive empathy measured using the ‘Reading the Mind in the Eyes’ test in a sample of more than 88,000 individuals of European ancestry ^10^. Here we draw several distinctions between the current study and the earlier study. First, the Triangle Task requires making inferences about mental states to animate non-social shapes, while the Eyes Test requires identifying the mental state from photographs of human eyes. Second, the current study investigates the genetic architecture of theory of mind at a specific age (13 years old). This allows for interrogation of the genetic contribution in adolescence when individuals are particularly vulnerable to several psychiatric conditions. However, we note that this is an age when the participants are undergoing puberty. Specific aspects of theory of mind develop differently during the course of puberty^48,49^. We were unable to account for differences in pubertal development in the current study.

In conclusion, this study does not find a significant genetic contribution to first-order theory of mind in adolescents. We find limited evidence that genetic variants that contribute to risk for psychiatric conditions predict variance in theory of mind ability in adolescents. However, we do find that genetic variants contributing to cognitive aptitude and cognitive empathy are significantly associated with theory of mind ability in adolescence. We speculate that observed differences in theory of mind in individuals with psychiatric conditions may be due to both biological and non-biological factors, or other biological phenotypes that mediate performance on tasks of theory of mind.

## Acknowledgements

This study was funded by grants from the Templeton World Charity Foundation, Inc., the UK Medical Research Council, the Wellcome Trust, and the Autism Research Trust. VW was funded by St. John’s College, Cambridge, and the Cambridge Trusts. The research was carried out in association with the National Institute for Health Research (NIHR) Collaboration for Leadership in Applied Health Research and Care East of England at Cambridgeshire and Peterborough NHS Foundation Trust. The views expressed are those of the author(s) and not necessarily those of the NHS, the NIHR or the Department of Health. We are grateful to all the families who took part in this study, the midwives for their help in recruiting them, and the whole ALSPAC team, which includes interviewers, computer and laboratory technicians, clerical workers, research scientists, volunteers, managers, receptionists and nurses. The UK Medical Research Council and Wellcome (Grant ref: 102215/2/13/2) and the University of Bristol provide core support for ALSPAC. GWAS data was generated by Sample Logistics and Genotyping Facilities at Wellcome Sanger Institute and LabCorp (Laboratory Corportation of America) using support from 23andMe. We would like to thank the research participants and employees of 23andMe for making this work possible. We specifically thank the following members of the 23andMe Research Team: Michelle Agee, Babak Alipanahi, Adam Auton, Robert K. Bell, Katarzyna Bryc, Sarah L. Elson, Pierre Fontanillas, Nicholas A. Furlotte, Karen E. Huber, Aaron Kleinman, Nadia K. Litterman, Jennifer C. McCreight, Matthew H. McIntyre, Joanna L. Mountain, Carrie A.M. Northover, Steven J. Pitts, J. Fah Sathirapongsasuti, Olga V. Sazonova, Janie F. Shelton, Suyash Shringarpure, Chao Tian, Joyce Y. Tung, Vladimir Vacic, and Catherine H. Wilson. This work was supported by the National Human Genome Research Institute of the National Institutes of Health (grant number R44HG006981). This publication is the work of the authors. Varun Warrier and Simon Baron-Cohen serve as guarantors for the contents of this paper.

### Additional information

The authors declare that they do not have any competing interests.

### Author contributions

VW co-designed the study, conducted the analysis and wrote the first draft. SBC co-designed the study, edited the manuscript, and obtained funding for the study.

